# Chromatin Reprogramming of In Vitro Fertilized and Somatic Cell Nuclear Transfer Bovine Embryos During Embryonic Genome Activation

**DOI:** 10.1101/2023.04.10.536281

**Authors:** Edward J. Grow, Ying Liu, Zhiqiang Fan, Iuri Viotti Perisse, Tayler Patrick, Misha Regouski, Sean Shadle, Irina Polejaeva, Kenneth L. White, Bradley R. Cairns

## Abstract

Reprogramming of the gamete into a developmentally competent embryo identity is a fundamental aspect of preimplantation development. One of the most important processes of this reprogramming is the transcriptional awakening during embryonic genome activation (EGA), which robustly occurs in fertilized embryos but is defective in most somatic cell nuclear transfer (SCNT) embryos. However, little is known about the genome-wide underlying chromatin landscape during EGA in SCNT embryos and how it differs from a fertilized embryo. By profiling open chromatin genome-wide in both types of bovine embryos, we find that SCNT embryos fail to reprogram a subset of the EGA gene targets that are normally activated in fertilized embryos. Importantly, a small number of transcription factor (TF) motifs explain most chromatin regions that fail to open in SCNT embryos suggesting that over-expression of a limited number of TFs may provide more robust reprogramming. One such TF, the zygotically-expressed bovine gene DUXC which is a homologue of EGA factors DUX/DUX4 in mouse/human, is alone capable of activating ∼84% of all EGA transcripts that fail to activate normally in SCNT embryos. Additionally, single-cell chromatin profiling revealed low intra-embryo heterogeneity but high inter-embryo heterogeneity in SCNT embryos and an uncoupling of cell division and open chromatin reprogramming during EGA. Surprisingly, our data also indicate that transcriptional defects may arise downstream of promoter chromatin opening in SCNT embryos, suggesting additional mechanistic insights into how and why transcription at EGA is dysregulated. We anticipate that our work will lead to altered SCNT protocols to increase the developmental competency of bovine SCNT embryos.

## Introduction

During the maternal-to-zygotic transition (MZT), embryos reprogram unipotent gamete chromatin into a totipotent state to allow for development^1,2^. Although it is known which loci in sperm and egg chromosomes are reprogrammed at the epigenome level in fertilized embryos^3^, it is still unclear how somatic cell nuclear transfer (SCNT) embryos also achieve reprogramming despite having a profoundly different initial starting epigenome. A critical aspect of this reprogramming is preparation and navigation of the transition from the transcriptionally quiescent state to a transcriptionally active state during the process of embryonic genome activation (EGA)^1,2^. EGA generates new gene products involved in transcription, splicing, chromatin regulation, and maternal mRNA clearance that help to complete MZT. However, transcriptional profiling comparing fertilized to SCNT embryos revealed that although EGA occurs in SCNT embryos, the specificity of which genes activate is profoundly dysregulated^4,5^.

A breakthrough in the mechanistic understanding of why most mouse SCNT embryos typically arrest at the EGA stage was the discovery that the repressive mark H3K9me3 from somatic cells acts as a barrier to proper EGA^4–6^. H3K9me3 decorates much of the somatic epigenome and is inherited and propagated in the first several cell divisions of SCNT embryos. Over-expression of the H3K9me2/3 demethylase (KDM4) in SCNT embryos removes H3K9me3, allows SCNT embryos to activate many critical EGA genes, and promotes development to the blastocyst stage^4,5^. Although initially demonstrated in mouse SCNT embryos^4^, this treatment has been extended to human^5^, non-human primate^7^, porcine^8^, and bovine^6^ cloning with similar results.

However, the post-implantation gestational success of SCNT embryos created with KDM4 overexpression is still extremely poor, precluding its wide-spread application to routine SCNT production. To explain the poor post-implantation development of KDM4-treated SCNT embryos, we considered several non-mutually exclusive explanations. First, it is possible that removal of H3K9me3 in SCNT embryos promotes a more normal EGA transcriptome compared to traditional SCNT embryos, but the level of activation or perdurance of some EGA expression is still perturbed. Second, it is possible that genome-wide H3K9me3 erasure in SCNT embryos leads to off-target demethylation of loci that typically retain H3K9me3 even in fertilized embryos, such as imprinted regions, transposons, centromeres, or telomeres. This “off-target” H3K9-demethylation could heritably alter the chromatin status of these loci which in aggregate deleteriously affects post-implantation development of SCNT embryos. Third, it is possible that although EGA is more robust in SCNT embryos after H3K9me3 removal, additional epigenetic marks exist that are not reprogrammed and these persist during the preimplantation stage, altering expression of post-implantation genes and development. Distinguishing between these three possibilities requires a more extensive understanding of the chromatin differences between fertilized and SCNT embryos, and how those differences affect the fidelity of EGA. In addition, it is still unclear which sequence-specific transcriptional activators fully accomplish EGA even in fertilized embryos^9^, let alone how they are dysregulated in SCNT embryos. Furthermore, a method that classifies embryonic SCNT chromatin and can distinguish properly-reprogrammed from improperly/incompletely reprogrammed SCNT embryos would have high utility pre-transfer embryo selection.

To address these challenges, we have taken a genome-wide approach to map the open chromatin landscape in fertilized bovine embryos and compare it to open chromatin found in SCNT embryos. We find that the highly dynamic open chromatin landscape is gradually reprogrammed in bovine SCNT embryos; by the morula stage, most surviving SCNT embryos display open chromatin profiles highly similar to those observed in fertilized embryos. EGA genes that are marked by open chromatin in fibroblasts are correlated with higher expression in SCNT embryos, but surprisingly, prior chromatin openness in fibroblasts is not sufficient for certain genes to be induced at EGA. Additionally, single-cell open chromatin profiling allows for the first time the identification of the minority of SCNT embryos with high reprogramming signatures. We also find maternal allele-specific bias in transcription and chromatin openness in *Bos taurus* x *Bos indicus* fertilized embryos at EGA. Remarkably, unbiased analyses of chromatin regions that fail to reprogram in SCNT embryos reveals a limited number of transcription factor motifs, including DUXC, which is the homologue of mouse/human DUX/DUX4. Functional validation in bovine fibroblasts indicates that over-expression of DUXC is sufficient to induce ∼84% of all EGA transcripts that fail to activate properly in SCNT embryos, identifying this transcription factor as a putative major gene-specific regulator of EGA that could reprogram SCNT embryos—potentially as an alternative to less-targeted epigenome-wide reprogramming strategies.

## Results

### Open chromatin profiling reveals distinct dynamics in SCNT preimplantation embryos

To compare the chromatin landscape of bovine preimplantation embryos generated from two disparate starting points—in vitro fertilized (IVF) embryos containing egg/sperm chromosomes or SCNT containing a fibroblast nucleus—we performed active transposition into active chromatin (ATAC)-seq^10^. For both embryo types, we analyzed multiple developmental stages: a stage before embryonic genome activation (EGA, 2-cell stage), peri-EGA (4- and 8-cell stage), and post-EGA (morula and blastocyst stage). First, we found that biological replicates of each embryo type and developmental stage clustered together (Fig 1a, Supplementary Figure 1a). However, open chromatin profiles from 2-cell and 4-cell stage SCNT embryos cluster more closely to the fibroblast donor, indicating that they retain chromatin signature of the somatic donor cell during the first several cell cycles (Fig 1a). Both IVF and SCNT embryos from the 8-cell stage (corresponding to the major wave of EGA^11^) cluster together, and are distinct from post-EGA developmental stages (Fig 1a).

**Figure 1:**
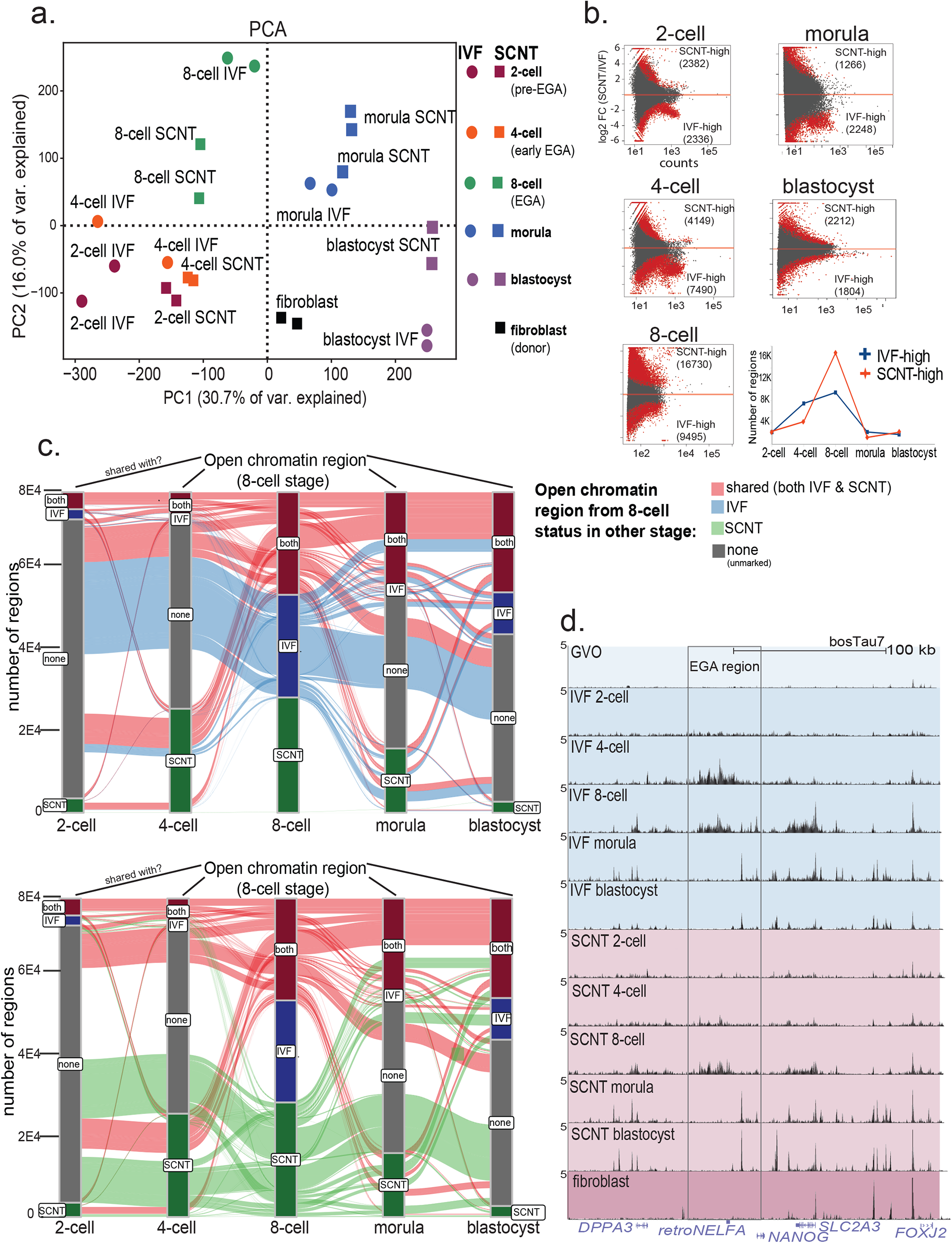
Global analysis of open chromatin dynamics in bovine IVF or SCNT preimplantation embryos. A. Principal component analysis of ATAC-seq data generated from pools of embryos at the indicated developmental stage. IVF embryos are depicted as circles, SCNT embryos are depicted as squares. Bovine fibroblasts are indicated as black squares. (n=2 biological replicates) B. Top: Differential open chromatin levels (ATAC-seq) in IVF vs SCNT embryos, each dot is a ATAC-seq peak region. Red dots indicate FDR<0.05 (DEseq2, n=2 biological replicates per each stage and embryo type). Bottom: quantification of the number of differential open regions. C. Alluvial plot showing developmental dynamics of ATAC-seq regions detected at the 8-cell stage in either IVF or SCNT embryos and whether they are marked as a peak in other indicated developmental stages. Peaks found in both IVF and SCNT embryos (shared) at the 8-cell stage are shown in red, IVF-only peaks at the 8-cell stage are shown in blue, SCNT-only peaks at the 8-cell stage are shown in green. (n=2 biological replicates per each stage and embryo type) D. Genome browser snapshots of ATAC-seq open chromatin data near the *DPPA3 (STELLA)/NANOG* locus showing developmental dynamics. Germinal vesicle oocyte data from (Pablo Ross, Nature Communications 2020). Tracks shown are the merged data from n=2 biological replicates per each stage and embryo type, y-scale=5 FPKM.

In order to quantify the number of loci that display differentially open chromatin levels, we compared IVF and SCNT embryos to each other at the same developmental stage. IVF and SCNT embryos at the 4-cell and 8-cell stage contained thousands of differentially open loci, with the 8-cell stage exhibiting the most difference in all developmental stages (Fig 1b). At the 8-cell stage, IVF embryos contained 9,495 regions with more open chromatin (“IVF-high”) while SCNT embryos contained 16,730 regions with higher levels of open chromatin (“SCNT-high”). Since this stage corresponds to the major wave of EGA in bovine embryos^11^, our chromatin profiling indicates global deviation of SCNT embryos from IVF embryos during the onset of transcription (Fig 1b). Interestingly, the post-EGA stages of morula and blastocyst had comparably fewer differentially open chromatin regions (Fig 1b), which we suspect is impacted by the survival bias of the minority of EGA-competent SCNT embryos while some EGA-incompetent SCNT embryos typically arrest by the 8-cell stage^6^.

Next, focusing on the 8-cell stage that exhibited the most different chromatin signature between IVF and SCNT embryos, we identified open chromatin dynamics across the developmental stages. We classified each region (n=81,000) at the 8-cell stage into IVF-specific, SCNT-specific, or shared. We found that 8-cell stage IVF embryos exhibited a striking pattern, in which most of the IVF-specific regions were not premarked (not open) even a cell-cycle earlier at the 4-cell stage and most of those regions were decommissioned (closed) by the morula stage 2 cell cycles later (Fig 1c). These results in bovine IVF embryos are consistent with human IVF embryos^12^. In contrast, many SCNT-specific regions were opened a cell cycle earlier at the 4-cell stage indicating a faster open chromatin establishment compared to IVF embryos (Fig 1c). Regions that were shared between IVF and SCNT embryos at the 8-cell stage were less dynamically developmentally regulated than SCNT-specific or IVF-specific regions, with ∼45% of these regions being retained in both morula and blastocysts stages (Fig 1c). Confirming the high level of developmental dynamics in chromatin landscapes, we found that the locus near the *DPPA3* (*STELLA*, a maternally-deposited transcript), *NANOG* (a component of the pluripotency network), and *NELFA* retrogene (induced at EGA) all showed distinct patterns across the stages sampled (Fig 1d). Of note, we noticed lower amounts of open chromatin at EGA (4-cell and 8-cell) stage in SCNT embryos compared to IVF embryos, consistent with the reported defects in EGA activation in SCNT embryos^6^ (Fig 1d).

### Dysregulation of open chromatin at EGA genes in SCNT embryos

Prompted by our observation that 8-cell stage embryos exhibited the most differences between IVF and SCNT embryos, we compared two genes that are highly induced at EGA. *ZSCAN4*, which is a conserved gene within mammals that functions to provide genome-stability^13^, exhibited higher levels of open chromatin in IVF embryos compared to SCNT embryos (Fig 2a, Supplementary Fig 2a). Consistent with its reported failure to activate properly at EGA in SCNT embryos, we observed lower *ZSCAN4* expression in SCNT embryos and the over-expression of the H3K9-demethylase KDM4E did not rescue this defect in SCNT embryos (Fig 2a). *SNAI1*, which encodes a transcriptional repressor SNAIL, also shows lower open chromatin at the 4-cell stage in SCNT embryos and this correlates with its failure to transcriptionally activate at EGA in SCNT embryos (Fig 2a). Interestingly, *SNAI1* is an actively transcribed gene in bovine fibroblasts and contains high levels of open chromatin at its promoter in fibroblasts (Fig 2a, Supplementary Fig 2a). Thus, open chromatin and active gene transcription in fibroblasts is not sufficient to confer gene activation in SCNT at EGA. We noticed that EGA genes (n=570) displayed more open chromatin at the promoter and directionally throughout the gene body extending beyond the annotated TES, consistent with prior reports in human and mouse embryos at EGA^3,12^ (Fig 2b). Consistent with the global lower levels of EGA expression in SCNT embryos, we found that on average, EGA genes in SCNT embryos failed to activate this signature of broad open-chromatin that extends through the gene body (Fig 2b).

**Figure 2:**
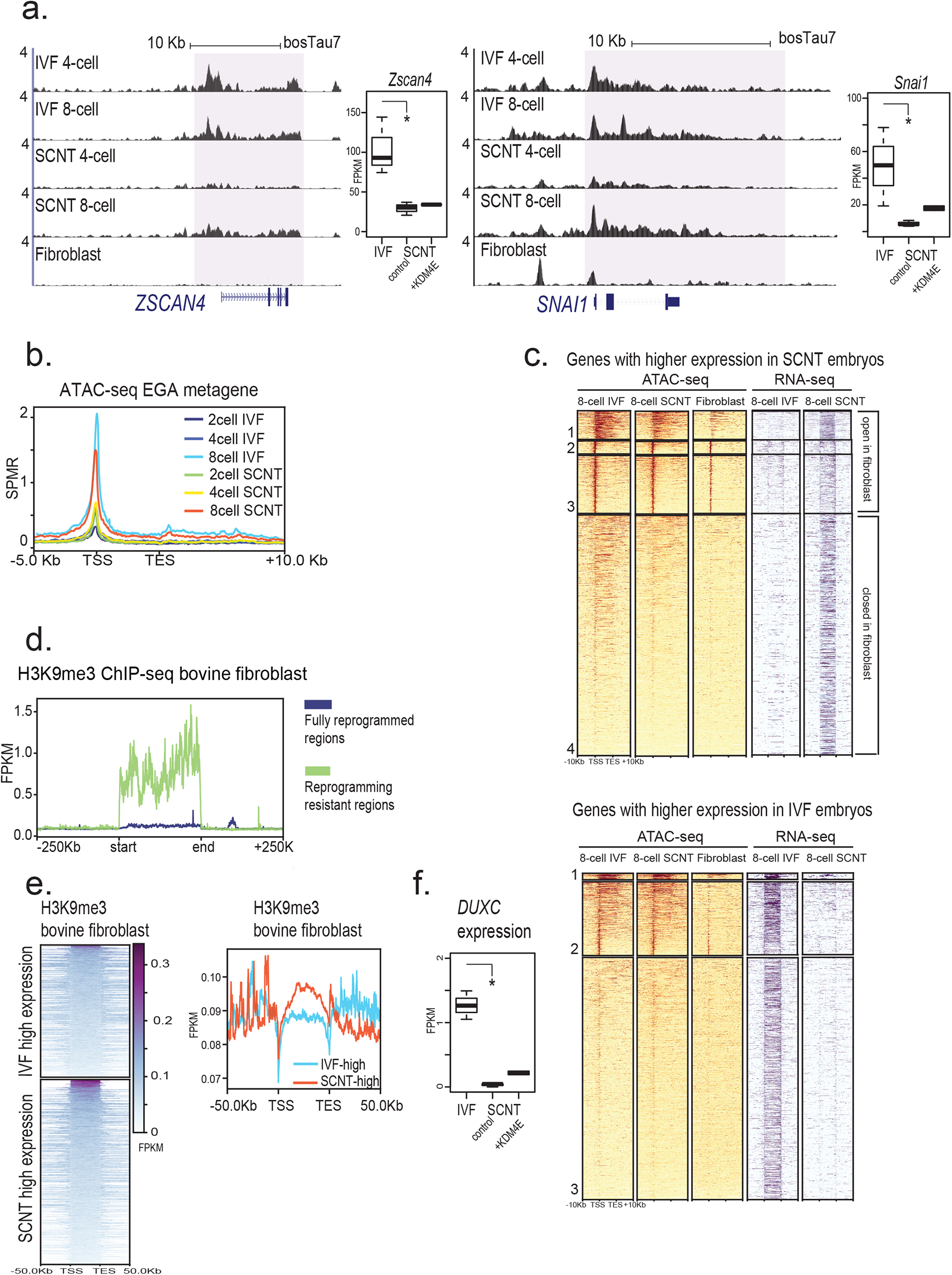
Chromatin reprogramming defects in SCNT embryos correlate with EGA dysregulation. A. Genome browser snapshots of EGA genes *ZSCAN4* and *SNAI1* (*SNAIL1*) showing lower levels of open chromatin in SCNT embryos compared to IVF embryos. ATAC-seq, n=2 biological replicates per each stage and embryo type, y-scale=4 FPKM. RNA-seq data reprocessed from Liu, et al. Development 2018, Significance is FDR<0.05, DEseq2, RNA-seq replicates n=3 biological replicates. B. Metagene profiles of EGA gene open chromatin showing broad open chromatin regions throughout the gene body preferentially in IVF compared to SCNT embryos. ATAC-seq, n=2 biological replicates per each stage and embryo type. C. Heatmap analysis of open chromatin levels at non-maternal EGA gene classes, IVF-high or SCNT-high, defined by Liu, et al. *Development* 2018. D. Metagene profile of H3K9me3 ChIP-seq from bovine fibroblasts (n=2 biological replicates) at Reprogramming Resistant Regions “RRRs” (IVF-high) or Fully reprogrammed regions EGA genes (defined by Liu, et al. Development 2018). E. Gene-level heatmap and metagene plot for indicated classes of genes (IVF-high or SCNT-high) using H3K9me3 ChIP-seq in bovine fibroblasts (n=2 biological replicates). F. Expression level of bovine *DUXC* transcript in 8-cell stage embryos: IVF, SCNT+KDM4E overexpression, or SCNT control showing higher expression of DUXC in IVF embryos. RNA-seq data from Liu, et al. Development, 2018, significance FDR<0.05 DEseq2.

Next, we compared the open chromatin levels at EGA genes that fail to activate properly in SCNT embryos (“IVF-high” genes) or were aberrantly high in SCNT embryos compared to IVF embryos (“SCNT-high” genes). Many of the SCNT-high genes were marked by open chromatin in the fibroblasts used as donors for SCNT (clusters 1, 2, 3 in Fig 2c), suggesting that in some instances, these genes might retain an epigenetic signature of the donor cell which persists during EGA in SCNT embryos. In contrast, most of the IVF-high genes were not marked by open chromatin in fibroblasts (Fig 2c), suggesting a need for their reprogramming in embryos. However, we still detected open chromatin in both IVF and SCNT embryos at IVF-high genes, indicating that open chromatin might be uncoupled from productive transcription at some sites in the genome during EGA, or potentially that chromatin opening precedes the accumulation of EGA transcripts.

Earlier work identified H3K9me3 in the epigenome of somatic cells that is thought to repress EGA gene expression in SCNT embryos and its removal by H3K9-demethylase (Kdm4 family) facilitates SCNT embryo EGA^4–6^. We confirmed this analysis in bovine SCNT embryos by comparing H3K9me3 levels from fibroblasts in what are termed “reprogramming resistant regions” or RRRs^4,5^ (Fig 2d). The mechanistic basis of the positive effect of KDM4-over-expression on EGA gene expression in SCNT embryos was thought to be a direct repressive effect on EGA genes. However, our reanalysis shows that only small fraction of either IVF-high or SCNT-high genes are substantially marked by H3K9me3 at either their promoter or coding region (Fig 2e). Additionally, reanalysis of published datasets indicates that the transcription factor DUXC (the homologue of the mouse and human Dux/DUX4 respectively, capable of strongly activating EGA genes expression in both of those species) is also expressed at very low levels in SCNT bovine embryos (Fig 2f). KDM4-overexpression in bovine SCNT embryos weakly reactivates DUXC expression, but it is possible that this level of DUXC is enough to activate DUXC-targets at EGA. Importantly, the majority of the effect size of KDM4E expression on mouse SCNT ZGA was mediated through DUX, as mouse SCNT embryos generated from DUX KO donor cells showed much lower development rates than mouse SCNT embryos over-expressing KDM4B using DUX WT donor cells^14^. Taken together, these data prompt a more nuanced explanation for the molecular mechanism through which KDM4 overexpression facilitates EGA in SCNT embryos (see Discussion).

### Single cell open chromatin profiling reveals inter-embryo heterogeneity and identifies the subset of well-reprogrammed SCNT embryos

In order to create high quality ATAC-seq maps, we initially pooled embryos from the same stage in order to obtain ∼100 cells, the lower limit for conventional ATAC-seq. However, we wondered if by pooling different embryos we might obscure the signature of reprogramming that exists in the minority of SCNT embryos that develop past the EGA stage. Single-cell ATACseq (scATAC-seq) is a robust method that allows for open chromatin profiling without the need for sample pooling^15^. We suspected that this approach might help distinguish between two possibilities: 1) that the lower levels of open chromatin at EGA genes in SCNT embryo pools is generated by the superposition of a minority of properly reprogrammed embryos combined with a majority of poorly reprogrammed SCNT embryos, or 2) all embryos and cells across SCNT embryos have a homogeneously low level of chromatin reprogramming. After producing 939 scATACseq libraries from either IVF or SCNT bovine embryos, we retained 819 high-quality scATAC datasets.

Open chromatin profiling by scATAC-seq reveals two main populations of cells: one that is enriched for earlier stage embryos and a minority population that is enriched for later stages (Fig 3a). In general, the separation between open chromatin states of IVF and SCNT embryos observed in pooled samples is also seen in scATACseq data (Fig 3a). Pseudotime analysis reveals that the number of cells in an embryo can be uncoupled from developmental pseudotime, however the minority of cells with late pseudotime are enriched for 8-cell stage and later which corresponds to the time of major EGA in bovine embryos^11^ (Fig 3b). Next, we quantified the open chromatin levels at all EGA genes, and as expected the later developmental pseudotime correlates with higher levels of open chromatin at EGA genes (Fig 3c). Some SCNT cells that exhibit high EGA open chromatin do not cluster with the late pseudotime cells, suggesting that genes that activate later in development after EGA or regions that inherit open chromatin from fibroblast donors that are not appropriately decommissioned likely also contribute to the developmental pseudotime score (Fig 3b). Importantly, the number of SCNT embryos that cluster with late developmental pseudotime is consistent with the proportion of SCNT embryos that progress past the EGA block at the 8-cell stage^6^ (8%, or 2/25 embryos).

**Figure 3:**
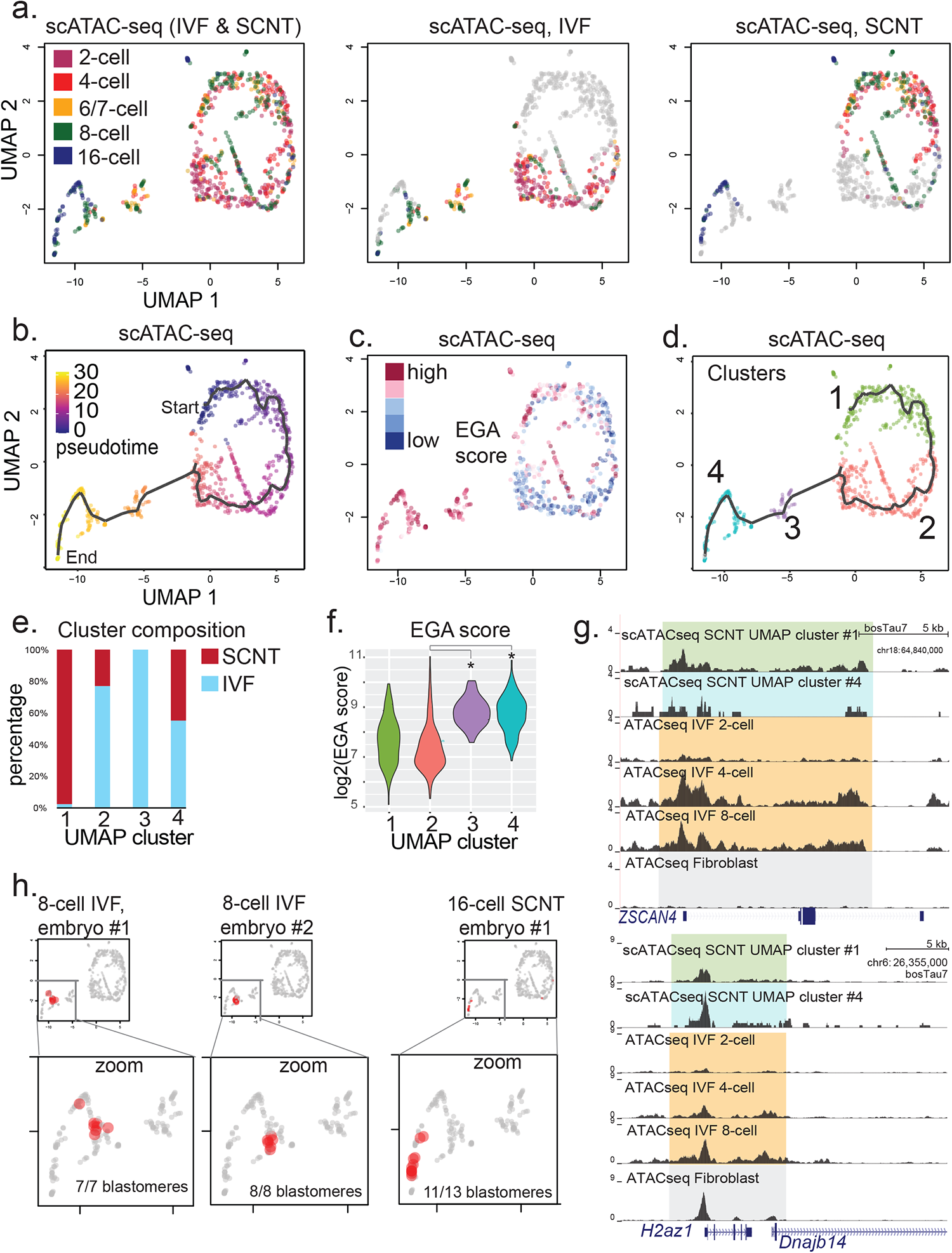
single-cell chromatin profiling reveals intra-embryo heterogeneity and EGA reprogramming signatures in a minority of SCNT embryos. A. UMAP analysis of scATAC-seq from indicated IVF or SCNT bovine embryos. Each dot is color-coded according to the cell number stage (left plot). IVF only cells (shown in middle) and SCNT only cells (shown on right). n=819 high-quality scATACseq libraries: n=73×2-cell SCNT, 118×4-cell SCNT, 165×8-cell SCNT, 64×16-cell SCNT, 147×2-cell IVF, 119×4-cell IVF, 117×8-cell IVF, and 1×16-cell IVF. B. Developmental Pseudotime analysis of scATAC-seq shows single, non-branching trajectory of early (“start”) to late development (“end”). C. UMAP analysis of scATAC-seq with indicated “EGA score”, which is derived from all non-maternal EGA genes in IVF embryos (n=533 regions, defined by Graf, et al. PNAS 2014). D. UMAP analysis of scATAC-seq with four cell clusters defined. E. UMAP composition of each cluster #1-4 indicating cluster #1 and cluster #3 are almost exclusively SCNT and IVF respectively, while clusters #2 and clusters #4 contain both IVF and SCNT cells. F. Quantification of the EGA score (IVF embryo defined EGA n=533 genes, from Graf, et al. PNAS 2014) at EGA n=5998 ATACseq peaks of each UMAP cluster #1-4. Significance is marked by Wilcox-Mann-Whitney test, FDR <0.05. G. Genome browser snapshots at *ZSCAN4* (IVF-high) or *H2AZ1* (properly reprogrammed) genes of in silico combined scATACseq data for clusters #1 or #4 compared to bulk ATAC-seq maps from IVF embryos at indicated stages. Bulk ATAC-seq, n=2 biological replicates per each stage and embryo type. H. UMAP plot of scATAC-seq comparing multiple blastomeres from the same embryo showing low intra-embryo heterogeneity.

Cluster analysis of the scATAC-seq data reveals four clusters of cells: clusters one and two are dominated by SCNT and IVF cells respectively, enriched for 2-cell and 4-cell stage embryos (Fig 3d). Cluster three is comprises IVF cells and cluster four contains both IVF and SCNT cells (Fig 3d). Cluster three and four correspond to later stage developmental pseudotime and exhibit higher open chromatin levels at EGA genes than clusters one and two (Fig 3e).

The scATAC-seq datasets allow us to combine *in silico* classes of cells without relying on developmental stage, which we found can be uncoupled from open chromatin reprogramming status (Fig 3a). We compared SCNT embryos from cluster one (lower EGA score) and cluster four (higher EGA score). Interestingly, for the *ZSCAN4* gene that fails to activate properly in SCNT embryos, the cluster four SCNT embryo cells have lower levels of open chromatin compared to 8-cell stage IVF embryos, suggesting that not just initial chromatin opening, but also the perdurance of chromatin opening might be critical for gene expression at EGA (Fig 3f). In contrast, the *H2AFZ* gene which encodes the histone variant H2AZ is open in fibroblasts and in both clusters one and four in SCNT embryos, consistent with its successful expression at EGA in SCNT embryos (Fig 3f). Since the chromatin landscapes are highly dynamic over time (Fig 1c), it is possible that SCNT embryos that continue to develop past the 8-cell stage have missed a “window of opportunity” in order to reprogram chromatin at critical EGA genes.

Since we manually dissociated embryos while retaining the identity of each cell in our dataset, we next asked if cells within the same embryo exhibit heterogeneity or homogeneity with their sister blastomeres. Remarkably, we found that on average, cells from the same embryo exhibited a highly similar open chromatin state, with their distribution along developmental pseudotime likely representing inter-embryo variation in developmental timing from the same batch of IVF or SCNT (Fig 3g). Low heterogeneity between blastomeres of the same embryo indicates that removing a single cell for a biopsy at these early stages may predict developmental success of the embryo at later timepoints. Taken together, our scATACseq dataset reveals high heterogeneity between embryos but low intra-embryo heterogeneity and reveals the minority signature of EGA-competent SCNT embryos. The upstream molecular determinants that enable a minority of SCNT embryos to show an EGA-competent chromatin signature is unclear, although we note similarities between the stochastic acquisition of ZGA/EGA transcriptome state in mouse two-cell like cells (2CLC)^16,17^, human 8-cell like cells (8CLC)^18,19^, or FSHD myoblast cultures^20^.

### Parent of origin allelic analysis reveals maternal contribution bias to EGA

Next, because parentally distinct gamete chromatin might uniquely affect early development in fertilized embryos (and could potentially be difficult to recapitulate in SCNT embryos), we asked whether allelic bias existed between the maternal or paternal allele at EGA in bovine IVF embryos. In order to generate F1 IVF bovine embryos, we used egg donors from pools of *Bos Taurus* cows and a single *Bos Indicus* bull. Whole genome-sequencing of egg donor pools or the single male parent combined with extensive filtering steps allowed us to identify approximately 6 million high-confidence homozygous polymporphic SNV in the *Bos Indicus* male parent that we can leverage to analyze allele-specific behavior at the chromatin or transcript level (see supplemental figure X). Although we cannot rule out the possibility that *Bos Indicus* sperm chromosomes behave differently than Bos Taurus sperm chromosomes in a *Bos Taurus* egg, we note that F1 hybrids are viable and fertile. The density of homozygous polymorphic SNVs in male parent was enough to analyze ∼35% of all EGA genes at the transcript level and 51% of all 8-cell stage open chromatin regions. We found that open chromatin in F1 fertilized embryos was markedly biased to the maternal allele (Fig 4a). Consistent with this, RNA-seq analysis of single F1 bovine IVF embryos revealed maternal bias at non-maternally deposited EGA transcripts, although we did detect some paternally-biased transcripts as well (Fig 4b).

**Figure 4:**
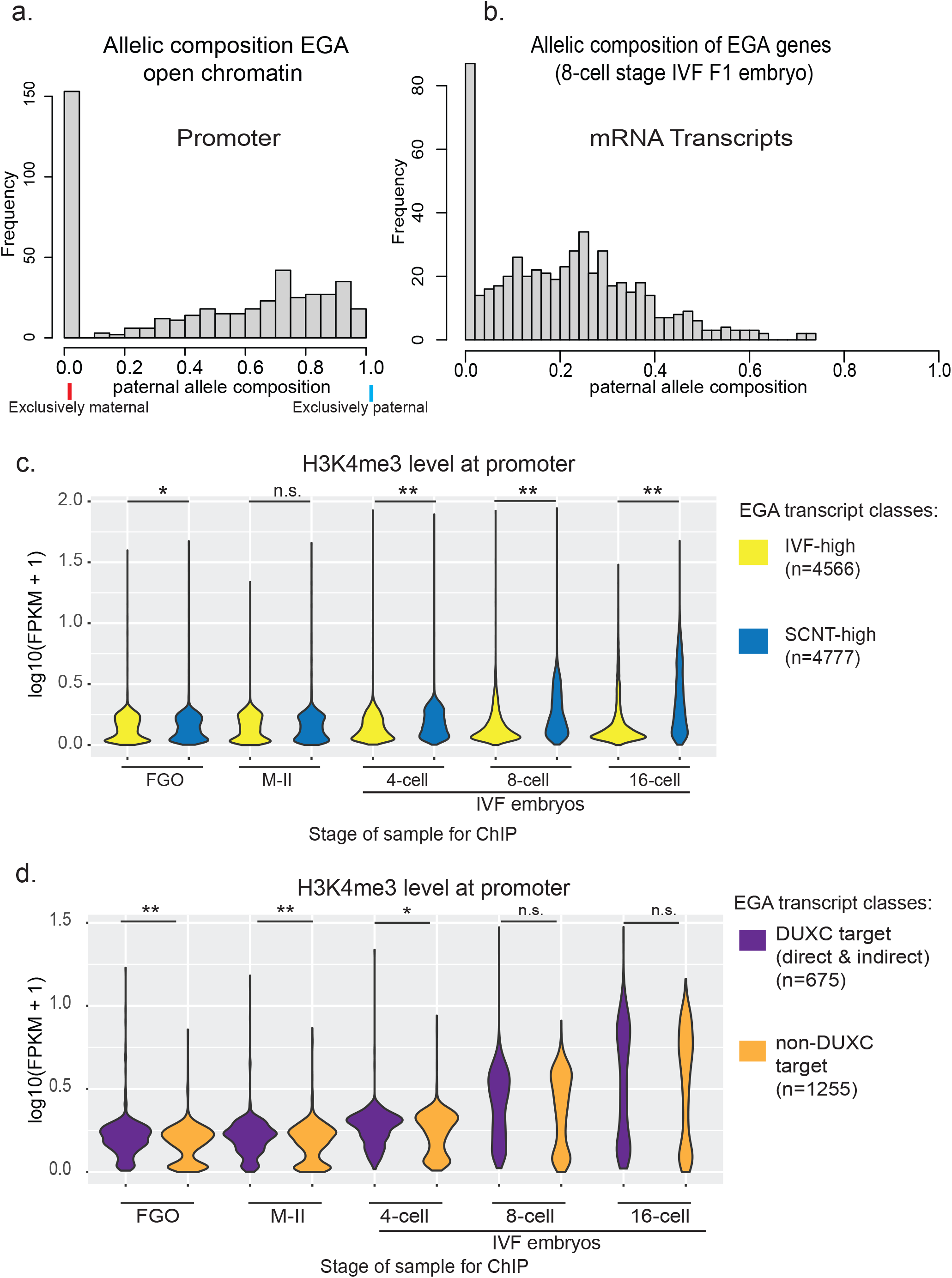
Allele-specific analysis of EGA chromatin and transcription in IVF embryos reveals maternal bias. A. Open chromatin allelic proportion of maternal vs paternal reads from F1 (*Bos Taurus* egg x *Bos Indicus* semen) ATAC-seq, restricted to ATAC-seq regions at the promoters of non-maternal EGA genes (n=533 regions, defined by Graf, et al. PNAS 2014). B. Expression analysis of allelic proportion of maternal vs paternal reads from F1 (*Bos Taurus* egg x *Bos Indicus* bull) RNA-seq 8-cell stage IVF embryos (n=8 single-embryo RNA-seq libraries). C. Integrative analysis of H3K4me3 ChIP-seq levels at indicated promoter classes in germinal vesicle oocytes, Metaphase-II eggs, or IVF embryos at indicated stages comparing two classes of genes: IVF-high or SCNT-high (RNA-seq from Liu, et al. Development 2018, H3K4me3 ChIP-seq from Lu, et al. *Science Advances* 2021). * denotes <0.05 FDR, * denotes <0.005 FDR, Wilcox-Mann-Whitney test. D. Integrative analysis of H3K4me3 ChIP-seq levels at indicated promoter classes in germinal vesicle oocytes, Metaphase-II eggs, or IVF embryos at indicated stages comparing two classes of genes: DUXC-targets (direct or indirect, genes that are induced after DUXC-overexpression in fibroblasts) or non-DUXC induced transcripts (not induced by DUXC-overexpression in fibroblasts), H3K4me3 ChIP-seq from Lu, et al. *Science Advances* 2021. * denotes <0.05 FDR, * denotes <0.005 FDR, Wilcox-Mann-Whitney test.

Next, we asked whether regions that showed differential open chromatin in SCNT vs IVF embryos were unusually marked in fertilized embryos. Reanalysis of H3K4me3 ChIP-seq data from bovine IVF embryos^21^ also revealed that SCNT-high EGA genes have higher levels of H3K4me3 in IVF embryos than IVF-high EGA genes (Fig 4c). The molecular basis of this phenomenon is unclear, but it suggests transcripts less sensitive to EGA defects in SCNT embryos (SCNT-high transcripts) might have an intrinsically stronger ability to recruit the H3K4me3 machinery during EGA in IVF embryos. Alternatively, these SCNT-high genes are potentially pre-marked by H3K4me3 in fibroblasts. In contrast, we found that DUXC targets (most of which are classified as IVF-high) are marked by higher levels of H3K4me3 in oocytes and eggs (but not at the 8-cell or 16-cell stage) compared to non-DUXC EGA targets (Fig 4d). Since DUXC targets are some of the highest and earliest expressed EGA genes^11^, this indicates that these genes might be pre-marked during oogenesis to allow for rapid activation (see discussion).

### Transcription factor contribution to open chromatin reprogramming in SCNT embryos

Open chromatin profiling has been used extensively to define the TF motifs present and to derive the master regulators of diverse cell states^15^. We anticipated that we could apply a similar bioinformatic approach to the open chromatin maps from IVF embryos and help define the EGA circuitry, and how this is perturbed in SCNT embryos. Importantly, our analysis of IVF embryos shows that a limited set of TF motifs dominate pre-EGA and EGA stages in bovine embryos (Fig 5a). These motifs include OTX2, KLF, and NFY motifs present at the 2-to-8-cell stage, and the DUX4 motif – which appears at the 4-cell stage (Fig 5a). Mechanistically, TF transcripts encoding KLF, OTX2, and NFY are maternally deposited while *DUXC* is zygotically expressed, likely explaining the emergence of DUX4 as a motif at the 4-to-8-cell stage, similar to human embryos^12^. These 4 motifs mark ∼80% of all open chromatin regions at the 8-cell stage, while other TF motifs such as GATA, OCT, and AP2 mark blastocyst stage IVF embryos which contain cell types such as trophectoderm and epiblast that are known to be regulated by these conserved TFs^9^ (Fig 5a).

**Figure 5:**
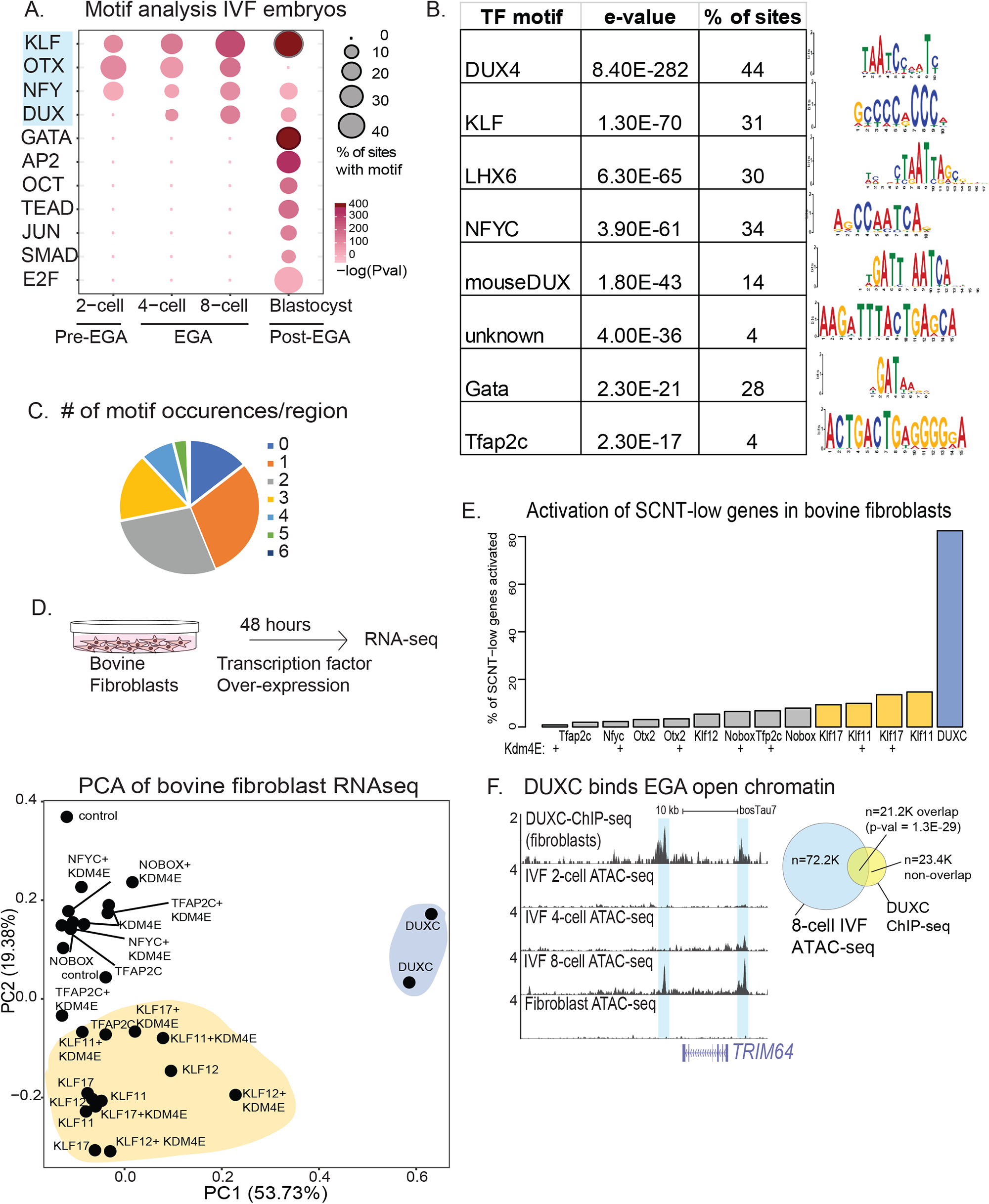
Motif analysis of putative TF regulators of EGA and SCNT reprogramming deficient chromatin regions. A. Transcription factor motif analysis of bovine IVF embryos at the indicated developmental stage (n=2 biological replicates per each stage). B. Table of TF motifs found at the SCNT-low open chromatin regions (FDR<0.05, DESeq2) from 8-cell stage. (n=2 biological replicates per each stage) C. Piechart of the number of individual motifs detected at each SCNT-low region, 8-cell stage embryos out of the top 8 motifs (n=2 biological replicates per each stage) D. Principal component analysis of RNA-seq analysis from bovine fibroblasts expressing indicated TF or TF+KDM4E combinations. (n=2 biological replicates per condition). Analysis using all expressed genes in the fibroblast dataset (EGA and non-EGA genes). E. Percentage of SCNT-low genes that are activated by indicated TF or TF+KDM4E over-expression in bovine fibroblasts. (n=2 biological replicates per condition) F. ChIP-seq of GFP-DUXC in bovine fibroblasts, genome browser snapshots showing EGA gene *Trim64* bound by DUXC at the regions of ATAC-seq open chromatin. (n=2 biological replicates per condition and stage, y-value = 4 FPKM for ATAC-seq, 2 FPKM for ChIP-seq). Right: Venn diagram of overlap between GFP-DUXC ChIP-seq data from bovine fibroblasts and ATAC-seq peaks in 8-cell stage IVF embryos, significance is measured using hypergeometric test with indicated p-value. G. TBD embryos experiment?

We reasoned that comparison of regions with significantly lower open chromatin levels from 8-cell stage SCNT embryos compared to 8-cell stage IVF embryos should reveal motifs that might underlie EGA defects in SCNT embryos. Motifs found at SCNT-low regions include DUX4, KLF, LHX6 (similar to homeobox OTX2), and NFY as the top enriched motifs (Fig 5b). Several additional motifs such as mouse DUX (partly divergent from human DUX4), GATA, or TFAP2C were also found but at a smaller percentage of SCNT-low sites (Fig 5b). Most of the SCNT-low sites (n=8,324) are marked by more than one of these motifs, 29.7% of sites have only one motif, and only 14% of sites have no match to the motifs on this list (Fig 5c).

Next, we tested whether the motifs identified in the open chromatin of SCNT-low regions might induce EGA genes that fail to activate in SCNT embryos. To conduct this experiment, we cloned candidate TFs (with expression evidence in IVF embryos) into transient expression vectors and co-delivered them into bovine fibroblasts with or without KDM4E overexpression. Here, we reasoned that some TFs may be sensitive to H3K9me3 that inhibits transcription from EGA from the somatic epigenome, while some TFs may be able to activate targets in fibroblasts regardless of H3K9me3 levels. Principal component analysis of the transcriptome bovine fibroblast RNA-seq samples show that ∼73% of the variance is explained by the first two components, with the first component (57.73% variance) dominated by the DUXC over-expression and the second component (19.4% variance) largely dominated by KLF factor over-expression (Fig 5d). Next, we restricted our analysis to only EGA transcripts that are “SCNT-low”. Surprisingly, we found that ∼84% of all SCNT-low EGA transcripts could be activated by DUXC overexpression, even though DUXC only activates ∼20% of all EGA transcripts, highlighting that the SCNT transcriptome at EGA is relatively similar to IVF embryos but specifically missing DUXC-target expression (Fig 5e). KLF11 or KLF17 also activated a minority (∼15%) of SCNT-low EGA genes, regardless of whether they were co-transfected with KDM4E. We note that KDM4E over-expression alone had minor effects on gene expression, indicating that although it can assist EGA activation in SCNT embryos, in fibroblasts it is not sufficient. This maybe be due to sequence-specific maternally deposited TFs that are present in embryos but absent in fibroblasts. Alternatively, the sensitivity of a certain TF to underlying H3K9me3 levels may be affected by dosage of the TF, and we emphasize that the levels of over-expression we achieve might overcome H3K9me3 repression. Importantly, one of the strongest and conserved targets of the DUX family of TFs is KDM4E (Supplementary figure X), which is strongly activated by DUXC overexpression and complicates the analysis of DUXC independence of H3K9me3 status.

### Bovine DUXC binds to EGA genes in fibroblasts and colocalizes with open chromatin regions from IVF embryos at EGA

Next, to determine the binding sites of bovine DUXC genome-wide, we performed ChIP-seq in bovine fibroblasts expressing GFP-DUXC. We find that 29% of all 8-cell stage open chromatin sites from IVF embryos are marked by a DUXC ChIP-seq peak, and this overlap is highly statistically significant (Fig 5f). DUXC, similar to mouse DUX and human DUX4^17^, does not require prior open chromatin in fibroblasts for its binding (Fig 5f), consistent with the properties of “pioneer” TFs. Future work will be required to test whether DUXC expression can facilitate EGA expression and development in bovine cloned embryos.

## Discussion

Inheritance of the somatic epigenome inhibits development and the precision of transcription program at EGA in SCNT embryos^4–7^. Prior work in several species indicate that inhibition of H3K9-methytransferases or over-expression of H3K9-demethylases is sufficient to promote preimplantation development and allows for a more normal transcriptome at EGA^4–7^. However, we reasoned that the low post-implantation development rate of these SCNT+Kdm4-OE embryos^4,6,7^ warranted a closer examination of the chromatin reprogramming deviation in SCNT embryos, with the goal of pinpointing TF(s) that could when overexpressed, reprogram the SCNT embryo during EGA without global off-target effects.

Here, we show that the open chromatin landscape is dramatically different in SCNT embryos pre- and peri-EGA but is mostly reprogrammed to match the IVF chromatin state by the morula stage in the minority of surviving SCNT embryos, 2 cell cycles later. Importantly, our data show that most open chromatin regions specific to the 8-cell stage IVF versus SCNT embryos are opened *de novo* (not detected by ATACseq in the 4-cell stage IVF embryo) and are decommissioned (closed) by the 32-cell morula stage just two cell cycles later—highlighting that SCNT embryos, in contrast to iPSC generation, have a narrow temporal window in which to reprogram. We also find that open chromatin in the fibroblast donor correlates with whether the SCNT embryos will be able to activate a particular gene at EGA, but also note that prior chromatin openness in the fibroblast donor is not sufficient to allow for proper activation at EGA in SCNT embryos. Our data show that chromatin opening is at some loci separable from the sequence specific TFs that either recruit RNA polymerase or promote productive transcriptional elongation—warranting closer investigation on the coordination of these two components of gene expression. Although we have focused on IVF-high open chromatin regions in 8-cell stage embryos, we note that the SCNT embryos also have *de novo* commissioning of SCNT-high regions during EGA and future work will be needed to characterize the genesis or TFs that drive this aberrant pattern.

Consistent with previous reports of “Reprogramming Resistant Regions” (RRRs)^4,5^, we find that SCNT-low EGA genes exist in Megabase-scale H3K9me3 regions. However, at the individual gene resolution in fibroblasts, there are only modest differences in H3K9me3 levels between IVF-high and SCNT-high genes. Complicating conclusions about H3K9me3 epistasis with EGA is the finding that DUXC is not activated properly in SCNT embryos but is weakly reactivated by KDM4-overexpression in SCNT embryos (Fig 2f), and one of the most evolutionarily consistent targets of DUX/DUX4/DUXC is the *Kdm4dl/KDM4E/KDM4E* gene (in mouse^17^, human^17^, and bovine respectively). Recent work has shown that most of the magnitude of the effect size on KDM4B-OE on promoting development of mouse SCNT embryos is eliminated by using *Dux* KO fibroblast donors^14^—raising the possibility that DUX factor expression is a key missing factor in mammalian SCNT EGA (see discussion below).

Our single-cell open chromatin approach has revealed the true signature of chromatin reprogramming in the minority of SCNT embryos that progress past the EGA-stage developmental block, and the high inter-embryo heterogeneity amongst SCNT embryos is consistent with ∼20% of these embryos developing to the blastocyst stage. Interestingly, we find heterogeneity in whether high-EGA score SCNT cells are assigned early or late developmental pseudotime, likely depending on additional parameter of the degree of fibroblast-specific chromatin decommissioning. Since we retained and monitored the individual embryo origin of all cells in our scATAC-seq dataset, we show for the first time that typically all cells within the same embryo show a homogeneous pattern of chromatin reprogramming versus non-reprogramming. Importantly, low intra-embryo heterogeneity indicates that single-blastomere biopsy of an early cleavage stage embryo could potentially predict the future developmental outcome. Certainly, a pre-selection of transferable embryos using a cell biopsy screening approach has been previously used extensively in cattle to control sex-selection of resulting offspring prior to embryo transfer to surrogate female recipient animals (Ken refs). A similar approach could be successfully used to identify a predictive epigenomic reprogramming signature to mitigate high gestational losses seen in clones. We envision that this could lead to term pregnancy success in SCNT embryos from edited fibroblasts that would facilitate its wide-spread adoption for the creation of genetically complex large animal models of human disease.

One of the outstanding questions in the SCNT reprogramming field is whether the parental-specific and unusual gamete epigenomes contribute to transcriptional defects at EGA during cloning, and if it is possible to modify the somatic cell donor to mimic this pattern. In support of this, we find that in DUXC targets have higher levels of H3K4me3 compared to non-DUXC EGA targets in oocytes and eggs (Fig 4d), suggesting that this subset of genes may be particularly set aside for pre-marking in the female gamete before EGA. However, DUX factors are not expressed in oogenesis and are instead zygotically activated. Future work will be required to understand how epigenetic marks are established in bovine oogenesis that prepare the chromatin template for EGA and how TFs interact with this gamete-specific landscape to accomplish transcriptional precision.

By using an F1 intercross between *Bos Taurus* and *Bos Indicus*, we find that there is a maternal allele bias in open chromatin regions at EGA, and analysis of allele-specific expression of EGA genes shows a similar maternal bias, consistent with transcriptional profiling of human uniparental embryos^22^. We emphasize that this result is not inconsistent with the report that the paternal chromosomes are highly transcribed during the earlier stage of minor ZGA^23,24^.

Analysis of our peri-EGA IVF embryo open chromatin reveals a small number of TF motifs: OTX2, KLF, NFY, DUX4, consistent with prior reports^9^. Importantly, the DUX4/DUXC motif is found at 44% of all chromatin regions that fail to open properly in SCNT embryos. Despite DUXC targets comprising only a minority of all EGA genes (∼20%) or open chromatin regions activated in IVF embryos, we nevertheless find that it can reactivate ∼84% of all SCNT-low EGA genes. By performing ChIP-seq with bovine DUXC in bovine fibroblasts, we find that DUXC possesses an almost identical DNA binding motif to human DUX4^17^. The intron-containing DUXC gene present in *laurasiatheria* is thought to be like the DUX progenitor gene that generated both the human DUX4 and mouse DUX intronless retrogenes^25,26^, confirming that outside of the rodent or primate clades that an ancestral DUX gene appears to be a key EGA factor in mammals. We emphasize that bovine DUXC is sufficient to activate SCNT-low genes in bovine fibroblasts and that the *DUXC* gene itself fails to activate normally in SCNT embryos, but we cannot rule out that there are other homeodomain TFs that may compensate for loss of bovine DUXC (like mouse *Obox4/Dux*^27^ or human *TPRX1/DUX4*^28^). Our findings raise the possibility that targeted TF expression in SCNT embryos may overcome EGA defects without non-targeted epigenome-wide heterochromatin erasure.

## Acknowledgements

We thank Kenneth Aston, Jingtao Guo, and members of the Cairns, Polejaeva, and White laboratories for fruitful discussions. We also thank B. Dalley in the HCI High-Throughput Genomics and Bioinformatic Analysis Shared Resource (NCI grant P30CA042014), J. Marvin and the University of Utah Flow Cytometry Facility (NIH, 1S10RR026802-01; NCI, 5P30CA042014-24). The NICHD (F32HD104442) to S.C.S., the NICHD (F32HD094500) and Lalor Foundation Fellowship 10041116 to E.J.G. and the Howard Hughes Medical Institute to B.R.C. and the NICHD (1R01HD095833) to B.R.C., I.P., and K.W.

## Material and Methods

### Pooled embryo ATAC-seq

All embryos were individually examined under a dissecting microscope to select embryos that had normal morphology with comparable sized blastomeres. For pooled samples, embryos were only pooled if they contained the correct number of blastomeres at the proper developmental timing. The zona pelucida was removed with 2mg/mL pronase at 37C for 2-3min. Polar bodies were removed if still present by gentle pipetting. Typically, for pooled sample ATAC-seq, we used around 100 total cells (50×2-cell embryos, 12×8-cell embryos, 3×32 cell stage morulas, or single blastocysts) into ∼1-1.5uL of PBS. Added 8.5 uL of ice-cold 1x lysis buffer (10mM Tris-pH=7.4, 10mM MgCl, 0.1% NP-40), pipette to resuspend, store on ice for 15 minutes to allow for cytoplasm to lyse. Add 12.5 uL of 2x Tris-DMF-tagmentation buffer (10mM Tris-pH=8.0, 5mM MgCl, 20% DMF). Add (batch-determined actitivty) 2 to 2.5 uL of Nextera Tn5 transposase (Cat# FC-121-1030, Illumina. Samples were incubated at 37C for 30 minutes in a thermomixer at 500 RPM shaking. The reactions were stopped with 0.5uL of 10% SDS, pipetted up and down, and placed on ice 5 min. 5x volume of PB buffer was added to the ATAC-seq reaction and purified using a mini-elute column (Qiagen Min**Elute** PCR purification Kit, Cat # 28006). Tagmentated DNA was eluted with 22uL of EB.

Libraries were PCR amplified in 50uL reaction volumes using custom Nextera PCR primers from Buenrostro, et al. Nature 2015. PCR reaction conditions: ∼20uL tagmented DNA, 25 uL of 2xphusion PCR mastermix (**Phusion** High-Fidelity PCR Master Mix with HF Buffer – 100 rxns, NEB, cat #M0531S), 0.625uL of 100uM F primer, 0.625uL of 100uM R primer. QS to 50uL total with H20. Do 5 cycles of “pre-amp” PCR. PCR cycling conditions: 72C incubation for 5 min. 5 cycles of the following: 95C for 30 sec, 72C for 90 sec. Remove 5uL of the “pre-PCR” reaction for qPCR. qPCR reaction conditions: 20uL qPCR reaction: 5uL of pre-amp PCR product, 10 ul of 2x syber (SsoAdvanced **SYBR** Green Supermix, Biorad catalog #1725275), 0.25uL of 100uM F primer, 0.25uL of 100uM R primer. Run 30-40 cycles of qPCR with the same cycling conditions as above. Select the additional number of PCR cycles to perform as per Buenrostro, et al. Nature Methods 2013. Ampure bead purification (Beckman Coulter, catalog # A63881) as per manufacturer’s instructions.

### Illumina sequencing

Individual libraries were normalized to 10 nM, and equal volumes were pooled in preparation for Illumina sequence analysis. Sequencing libraries (25 pM) were chemically denatured and applied to an Illumina HiSeq paired-end flow cell with an Illumina cBot. Flow cells were then transferred to an Illumina HiSeq 2000 instrument and sequenced in 150-bp paired-end mode.

### Single-cell ATAC-seq

Embryos were processed (as described above for pooled samples). Each embryo was dissociated into single blastomeres in a single drop of Ca+2/Mg+2 free PBS+0.1% PVP-40 with gentle pipetting. Each individual blastomere of the embryo was pipetted into a PCR strip tube for lysis and ATAC-seq, retaining the embryo identity of each single-cell ATACseq library.

### Bovine fibroblast ATAC-seq

Bovine fetal fibroblasts (“Crushertime” bull) were processed for ATAC-seq as Hendrickson, et al. *Nature Genetics* 2017.

### Culture of bovine fibroblasts

Cells were maintained in DMEM-high glucose (Thermo-Fisher 11965118) + 10% FBS, 1X Non-essential amino acids, 1X glutamax, 1X penicillin/streptomycin at 20% oxygen, and passaged using trypsin every 2-5 days. For GFP-DUXC ChIP-seq experiments, cells were transfected with a piggy-back GFP-bovineDUXC plasmid or a GFP-control plasmid as a negative control.

### ChIP-seq of H3K9me3 and GFP-DUXC in bovine fibroblasts

Antibodies to H3K9me3 (active motif catalog #39161) and anti-GFP (abcam catalog #13970) were used for IP. For negative control, GFP-alone control BFF’s were used. ChIP was performed as described in Rada-Iglesias, *Nature* 2010. Cells were cross-linked with 1% formaldehyde for 10 min before being lysed. Chromatin was sonicated with a BioRuptor system (Diagenode). Cellular debris was pelleted, and the DNA was precipitated overnight at 4 °C. After reversal of cross-links, libraries were prepared with an NEBnext DNA Library Prep kit (NEB, E7370L). Adaptor-ligated DNA was size-selected and purified with AMPure XP beads (Beckman Coulter, A63881) before PCR using NEB 2xphusion PCR mastermix (NEB, cat #M0531S).

### RNA-seq from Bovine fibroblasts

RNA from bovine fibroblasts expressing GFP-DUXC or control plasmid was purified using Trizol (Thermo catalog #15596018). PolyA mRNA was enriched using NEB E7490S kit. RNA was fragment using NEB E6150S kit. RNA-seq libraries were constructed using NEB E7760S kit.

### RNA-seq from Bovine fibroblasts or F1 (Bos Taurus x Bos Indicus) IVF embryos

FACS sorted fibroblasts were extracted with Trizol to purify RNA. IVF embryos were lysed in lysis buffer as described and cDNA was prepared using SMART-Seq® v4 Ultra® Low Input RNA Kit for Sequencing (Takara Bio, catalong #634890). cDNA was tagmented using Nextera XT DNA Library Preparation Kit (Illumina catalog #FC-131-1024) and libraries were amplified by PCR as per ATAC-seq libraries above.

### RNA-seq analysis

RNA-seq reads were trimmed and filtered for quality using BBDuk and FastQC (version 0.10.1). Processed reads were aligned using TopHat2 version 2.1.0 (--t --q --N1 --L 25 --X 2000 --no-mixed --no-discordant), and counts for each transcript were generated using ‘Get datasets’ (https://metacpan.org/pod/distribution/Bio-ToolBox/scripts/get_datasets.pl). De novo transcriptome was assembled using all embryo and fibroblast RNA seq data with Stringtie version 1.3.6.

### ATAC-seq and ChIP-seq alignment and peak calling

Processed reads were aligned to bostau7 with Bowtie2 (v2.2.6) with the following parameters: (--t --q --N1 --L 25 --X 2000 --no-mixed --no-discordant). PCR duplicates were removed using prior to peaks calling (over input DNA for ChIP-seq) with MACS2 ‘callpeak’ (--f BAMPE --B -- SPMR).

### Differential expression or differential chromatin open analysis

For differential ATAC-seq analysis: peaks were called using MACS2 peak caller. Summits of regions were extended to 1Kb windows. Bedtools was used to intersect peaks regions from biological replicates to identify intersect regions and merge peak regions from all embryo and fibroblast samples into a union set. ATAC-seq reads were counted and assigned to regions and differential expression analysis was performed using DESeq2 (version 3.11), with FDR<0.05 being set as the significance cut-off.

### Overlap between ATAC-seq peaks between samples

Bedtools version 2.17.0 was used to identify overlap ATAC-seq or ChIP-seq peaks (called previously with MACS2).

### Alluvial plots

For each ATAC-seq dataset for each developmental time point and each type of embryo (SCNT vs IVF) regions were classified as “detected” if the independent MACS2 peak-calling of the biological replicates both called the region overlapping with a peak. Across the union set of ∼200,000 ATAC-seq regions (non-redundant union list of peaks from ATAC-seq experiments from embryo and fibroblasts), each region was classified as “IVF”, “SCNT”, “both”, or “none” if the region was found in IVF-only, SCNT-only, both IVF and SCNT, or neither IVF and SCNT samples respectively. In figure 1C, only ∼80,000 peaks that were found in 8-cell stage embryos (IVF or SCNT) were used. Alluvial plots were generated using the R plugin “alluvial”.

### Metagene and heatmap plots

Deeptools 2.0 package was used to create metagene plots and heatmaps for ChIP-seq/ATAC-seq data.

### UMAP analysis

Was conducted using R plugin “Monocle 3”. Briefly, single-cell ATAC-seq data were filtered for regions that were detected in >10 samples (out of n=819 scATAC-seq libraries), leaving n=319,000 regions. Number of dimensions was n=50, Clusters from UMAP analysis were identified using “resolution=2e-3”, and developmental pseudotime was calculated using monocle 3. After assigning scATAC-seq samples to clusters (n=4), samples were combined to create *in silico* bulk ATACseq.

### EGA score for UMAP depiction of scATAC-seq data

EGA transcripts (not expressed in eggs, FPKM<1, and induced in 8-cell and 16-cell stage IVF embryos, data from Graf, et al. PNAS 2014) n=4600 were selected. Then, we intersected these genes with the union set of ATAC-seq peaks (n=319,000) and were left with n=5998 ATAC-seq peaks. These peaks represent both promoter and gene body open chromatin, the latter of which is a strong feature of the EGA open chromatin landscape. For each single cell, reads were counted from each EGA gene-linked ATACseq peak and then normalized by the number of ATAC-seq peaks found in peaks (to account for sequencing depth and variation in signal captured from each single cell).

### Principal component analysis

GGplot2 was used for principal component analysis. A union-set of all summit-centered peaks (called from MACS2) of all ATAC-seq regions (detected in embryos and fibroblasts) was used and normalized read counts from each replicate of ATAC-seq library was calculated.

### Variant calling

Whole genome sequencing of the bovine ‘Crushertime’ fibroblast, *Bos Indicus* bull “Dream Boy”, and a pool of cumulus cells from egg donors (*Bos Taurus*) were used to with VarScan.v2.3.7 to identify homozygous SNVs using command *“--min-coverage 20 --min-var-freq 0.1 --p-value 0.05*”. Then, homozygous SNVs that were polymorphic between the male parent (*Bos Indicus*) and female parent (*Bos Taurus*) were detected in the F1 IVF embryos (cross used for scATAC-seq experiments) to calculate paternal allele expression. We also produced RNA-seq from *Bos Taurus* parthenote embryos, performed similar variant calling on them and filtered out any SNVs that were detected in *Bos Taurus* parthenotes and *Bos Taurus* x *Bos Indicus* IVF embryos. Reads that captured nucleotides that were identified as polymporphic, homozygous SNVs in Dream Boy but had no SNV were classified as maternal-allele expression. For our analysis of allelic bias of EGA genes (either through ATAC-seq in F1 or in RNA-seq in F1s), we only used EGA genes that were induced at EGA but not expressed in eggs or GV oocytes (FPKM<0.1) to avoid erroneously identifying allelic-expression as maternally biased when in fact, the transcripts were maternally-deposited.

### Reprocessed public datasets

RNA-seq from bovine IVF vs SCNT 8-cell stage embryos (Liu, et al. *Development*, 2018): GSE99294. RNAseq from bovine preimplantation embryos (Graf, et al. *PNAS* 2014): GSE52415. Bovine GVO ATAC-seq (Halstead, et al. *Nature Communications*, 2020): GSE143658. H3K4me3 Cut-N-Run from bovine oocytes/eggs/IVF-embryos (Lu, et al. *Science Advances,* 2021), GSE163620.

### Motif analysis

After peak calling with MACS2, summits were identified and windows were extended 250bp from the summit nucleotide. MEME-ChIP suite (Version 5.5.1) was used: Philip Machanick and Timothy L. Bailey, “MEME-ChIP: motif analysis of large DNA datasets”, *Bioinformatics* 27(12):1696-1697, 2011.

### Oocyte collection and in vitro maturation

Bovine ovaries were collected from a local abattoir (JBS, Hyrum, Utah) and transported in a cooler containing 0.9% saline solution to the laboratory. Because this research used discarded tissues to generate embryos that were then disposed of without implantation, approval of this work via the Institutional Animal Care and Use Committee was not required. The cumulus-oocyte complexes (COCs) were aspirated from 3-8 mm follicles using an 18-gauge needle and vacuum system. Only compact COCs with homogenous ooplasm and intact layers of cumulus cells were used for somatic cell nuclear transfer (SCNT) or in vitro fertilization (IVF). Following aspiration, COCs were cultured at 38.5°C with 5% CO_2_ for 22–24 h. The oocytes were cultured in TCM199 maturation medium containing Earles salts, L-glutamine, and sodium bicarbonate (Sigma M4530) supplemented with 10% (v/v) fetal bovine serum (FBS) (Hyclone, SH30070.03), 0.05 mg/ml of bovine follicle stimulating hormone (Sioux Biochemicals, Sioux city, IA), 5 mg/ml of bovine luteinizing hormone (Sioux Biochemicals, Sioux Center, Iowa), 100 U/ml of penicillin, and 100 mg/ml of streptomycin (Gibco 15070-063).

### In vitro fertilization

Following 22–24 h of maturation, MII oocytes were fertilized by using the laboratory’s standard in vitro fertilization protocol [Reproduction, 2006, 131:45-51]. Briefly, one straw of cryopreserved bovine semen (Hoffman AI, Logan, Utah) was removed from the liquid nitrogen tank and placed into a 35°C water bath to thaw. Live sperm were isolated by centrifugation through a 45%/90% percoll gradient (MP Biomedical, Irvine, California), suspended (final concentration 1×10^6^/ml) in Tyrodes albumin lactate pyruvate containing 10 μg/ml porcine heparin and used to fertilize the oocytes. Twenty to 22 h post-IVF, cumulus cells were removed by vortexing, and fertilized zygotes were washed in phosphate buffered saline with 0.32 mM sodium pyruvate, 5.55 mM glucose, and 3 mg/ml bovine serum albumin (PB1+). After washing, zygotes were cultured in 40 μl of synthetic oviductal fluid (SOFaa) [Human Reproduction 2000; 15:395-401] with 3% FBS (Hyclone, SH30070.03, Logan, Utah) overlaid with mineral oil. The embryos were cultured at 38.5°C in a humidified atmosphere with 5% CO_2_. Half of the SOFaa medium was removed and replaced with fresh, equilibrated medium every other day starting the day after in vitro culture. Historically, production of IVF embryos via these methods followed by blastocyst embryo transfers to cattle recipients has generated successful pregnancies at an approximate rate of 50% (number live births/number of embryo transfer procedures) by our research group (unpublished observations). The timing of embryo collection post-fertilization was approximately 24–36 h for 2-cell stage, 48–60 h for 8-cell stage, 156–168 h for morula stage, and 192–204 h for blastocyst stage. Embryos that were obviously dead or arrested by microscopic examination were removed from culture.

### Somatic cell nuclear transfer (SCNT)

Primary bovine fibroblast cultures were established from ear biopsy tissues from multiple donor Brahma Spanish bull crosses by using well-established procedures [PNAS USA; 2000; 97:990-995]. These cell lines have been previously employed on multiple projects to evaluate their efficiency to generate live calves via SCNT (unpublished observations). Frozen/thawed cells were grown to 80-100% confluence then passaged, with cells from passages 4-5 used as nuclear donors. Three days prior to a cloning session, donor fibroblast cells were thawed and propagated, and incubated in DMEM medium (Hyclone, Logan, Utah) supplemented with 15% (v/v) FBS at 39°C with 5% CO_2_. Bovine fibroblasts at 80-90% confluence were serum-starved by replacing culture medium with DMEM containing 0.5% (v/v) FBS 24 h prior to SCNT. Oocytes were matured for 18–20 h, then denuded by using 100 μl of 1% (v/v) hyaluronidase (Sigma H3506), incubated for 5 min at 38.5°C with 5% CO_2_, followed by gentle pipetting. The denuded oocytes were then rinsed with PB1+, and oocytes with polar bodies were selected and separated. SCNT was performed according to established protocols [Reproduction 2006; 131:45-51; Anim Reprod Sci 2006; 95:234-243; Mol Reprod Dev 2004; 68189-197]. Briefly, oocytes with polar bodies were incubated in 0.6 μg/ml demecolcine for 30–40 min and a metaphase plate and a polar body were removed by using a beveled pipette. Fibroblast cells were covered in 0.25% trypsin (Gibco, Carlsbad, CA) for 1 min, and then the trypsin was removed, and fibroblasts were incubated for another 6 min at 39°C with 5% CO_2_. Once fibroblast cells were detached, they were rinsed in warm medium and centrifuged at 150 × *g* for 6 min. The supernatant was removed, and the cellular pellet was resuspended in 100 μl of Hepes SOF medium [Hum Reprod 2000; 15:395-401]. One fibroblast cell was injected into the perivitelline space of the recipient oocyte and fused by using one direct pulse of 1.2 kV/cm for 22 μs by an Electro Cell Manipulator 2001 (BTX, San Diego, CA) in 0.28 M sorbitol, 0.05% (w/v) BSA, 0.1 mM CaCl_2_, 0.5 mM MgCl_2_, and 0.5 mM Hepes. Following fusion, embryos were washed through Hepes SOF, and incubated in embryo culture medium for 1 h. Activation was then performed, with successfully fused embryos cultured in 5 μM ionomycin for 5 min, followed by 4 h of incubation in activation medium composed of SOFaa medium with 10 μg/ml cyclohexamide and 1 mM 6-dimethylaminopyridine at 38.5°C with 5% CO_2_. Following activation, embryos were cultured in 40 μl drops of SOFaa with 3% (v/v) FBS overlaid with mineral oil. The embryos were cultured at 38.5°C in a humidified atmosphere with 5% CO_2_. Half of the SOFaa medium was removed and replaced with fresh, equilibrated medium every other day starting the day after in vitro culture. The timing of embryo collection was the same as for IVF embryos as described above. The rates of development for IVF and SCNT embryos were similar at each developmental stage.

### Bovine Fetal fibroblast (BFF) isolation

BFF cells were isolated from a male day 55 fetus as previously described and cultured in Dulbecco’s modified Eagle’s medium ((DMEM) high glucose; Cytiva, Logan, UT) supplemented with 15% of fetal bovine serum (FBS; Cytiva, Logan, UT) and 100 U/mL penicillin-streptomycin (Life Technologies) at 38.5°C in an atmosphere of 5% CO2 in air^29^ .

### Transfection of BFF cells

First, we transfected BFF cells with 23 combinations of constructed vectors containing either GFP control or KDM4e-GFP along with mCherry ligated to a transcription factor (TF). We tested the following TFs: OTX2, NFYC, Tfap2c, Cdk9, DUXC-DEL, NOBOX, KLF11, KLF17, and KLF12. In addition, we transfected cells with either mCherry control or mCherry-KDM4e with GFP-DUXC, and transfection fluorescence control samples containing GFP-KDM4e and mCherry control; mCherry-KDM4e and GFP control; and GFP and mCherry controls. After analysis of the mRNA of the cells, we performed additional transfection with the following combinations: KLF11/KLF12/KLF17/DUXC/NOBOX, KLF11/KLF12/KLF17/DUXC/NFYC, KLF11/KLF12/KLF17/DUXC/Tfap2c, KLF11/KLF12/KLF17/DUXC/GFP, and GFP and mCherry controls. All transfections were performed using the Lonza Amaxa 4D-Nucleofector System (Program EH-100) in a 100µl nucleovette system following the manufacture’s instructions. We transfected 2.8×10^6^ BFF cell that were cultured in T-25 flask until confluence and after transfection, the cells were placed in a single well of a 6-well plate for recovery. Past 24h post-transfection, cells had the GFP and mCherry fluorescence checked under the microscope and 48h hours after the transfection the cells were submitted to Fluorescence-Activated Cell Sorting.

### Fluorescence-Activated Cell Sorting (FACS)

Individual transfected cells were sorted using a FACSAria II (BD Bioscience) setup with a 70 nm nozzle and using FACSDiva software (version 6.1.3) with the “purity” sorting setting. Forward scatter (FSC) and side scatter (SSC) were used to identify cells and eliminate doublets. GFP and mCherry fluorescence were measured using a blue excitation laser (488 nm) with detection in the GFP channel (530/30 filter) and a yellow green laser (561 nm) with detection in the mCherry channel (610/20 filter), respectively. Cells were sorted for double-positive GFP and mCherry fluorescence with thresholds determined based on negative controls. The double-positive selected cells were spun and resuspended in 500 µl of TRIzol (Invitrogen) and immediately placed into the -80 C freezer for further RNA-sequence analysis.

